# Effects of hM4Di activation in CamKII basolateral amygdala neurons and CNO treatment on Sensory-Specific vs. General-PIT; refining PIT circuits and considerations for using CNO

**DOI:** 10.1101/700120

**Authors:** Rifka C. Derman, Caroline E. Bass, Carrie R. Ferrario

## Abstract

Pavlovian stimuli can influence instrumental behaviors, a phenomenon known as Pavlovian-to-instrumental transfer (PIT). PIT arises via psychologically and neurobiologically independent processes as Sensory-Specific-PIT (SS-PIT) and General-PIT. SS-, but not General-PIT, relies on the basolateral amygdala (BLA), however the specific BLA neuronal populations involved are unknown. Therefore, here we determined the contribution of glutamatergic BLA neurons to SS-PIT. The BLA was transduced with virus containing either GFP or hM4Di, driven by the CamKII promoter. Rats were then tested for SS- and General-PIT following Vehicle or Clozapine-n-oxide (CNO, the hM4Di-activating ligand) injection. CNO had no effect on SS-PIT in the GFP control group, but selectively blocked its expression in the hM4Di-expressing group. Furthermore, CNO did not alter the expression of Pavlovian outcome devaluation effects in GFP or hM4Di expressing groups, indicating that the hM4Di-mediated loss of SS-PIT did not result from an inability to recall the sensory-specific details of the Pavlovian stimulus-outcome associations. Unexpectedly, CNO disrupted General-PIT in both GFP and hM4Di expressing groups, indicating that CNO alone is sufficient to disrupt affective, but not sensory-specific processes. Disruption of General-PIT by CNO was not due to generalized motor effects, but instead may be related to shifts in internal state produced by CNO. Together these data identify BLA CamKII neurons as critical for the expression of SS-PIT, and reveal important considerations for using CNO to study general affective motivation.

## I. INTRODUCTION

During Pavlovian appetitive conditioning, repeated pairing of a conditioned stimulus (CS) with a rewarding outcome results in the formation of an association between the stimulus and outcome (S-O). The formation of this association can be inferred through the manifestation of complex behaviors including the emergence of anticipatory conditioned responding directed toward the site of expected reward delivery (e.g., food cup approach). The nature of the S-O representation is composed of multiple distinct elemental associations between the CS and various experiential features of the outcome, including the outcomes sensory properties and a wide range of post-ingestive effects (e.g., hedonic, emotional, satiety, etc.; Konorski, 1967; Delamater, 2012). Importantly, these complex S-O associations can acquire the capacity to spontaneously modulate the expression of learned instrumental behaviors supported by response-outcome associations (R-O). This phenomenon, known as Pavlovian-to-instrumental transfer (PIT), is thought to play a role in a wide range of naturally occurring behaviors in both human and non-human animals, and may also contribute to the development of motivational disorders like addictions and obesity (Boutelle Bouton, 2015; Derman Ferrario, 2018; Watson et al., 2018).

In the laboratory setting, appetitive PIT can be measured by determining how the presentation of a previously established CS augments instrumental responding for the same or similar outcome predicted by the CS. Since its first demonstration by Walker (1942), PIT has been studied using a variety of paradigms, most notably, Single-Outcome PIT (SO-PIT), General-PIT, and Sensory-Specific PIT (SS-PIT). While not initially recognized as distinct, it is now established that these variants of PIT are psychologically and neuronally dissociable (Cartoni et al., 2016, for review). SO-PIT is observed when presentation of a learned CS excites instrumental responding for the same outcome predicted by the CS; critically in this procedure only one CS-outcome and one response-outcome association is trained (first demonstrated by Walker, 1942). General-PIT in contrast, is observed when CS presentation non-selectively excites instrumental responding for any outcome with similar general motivational properties or modality (i.e., ingestive, sexual, etc.), independent of the specific outcome associated with the instrumental response (first demonstrated by Balleine, 1994). Finally, SS-PIT is classically observed when presentation of a CS selectively excites instrumental responding for the same outcome predicted by that CS, as compared to instrumental responding for a different outcome (first demonstrated by Colwill Motzkin, 1994).

Of the three established forms of PIT, General- and SS-PIT are most strongly dissociable psychologically and neurobiologically (Cartoni et al., 2016). Sensory-specific associations carry information about the distinct sensory components of an experience, independent of affective influences; for instance, the flavor, viscosity, and temperature of an imbibed liquid, but not the general satisfaction, or relaxation that could result from its ingestion. Here, subsequent behaviors are influenced via activation of memories of these specific sensory properties (i.e., sensory-specific memories). In contrast, general affective associations carry information about the emotional content of an experience; for instance, the feeling of comfort associated with consuming a warm beverage on a cold day. Via the general affective mechanism, behaviors are influenced by memories of the emotional experience and general state.

Of course, a given experience can have both sensory-specific and affective properties. However, distinguishing between affective and sensory-specific processes, as can be achieved experimentally by examining SS- and General-PIT, has important and broad reaching implications for the study of motivated behaviors and associated diseases. For example, alterations in these Pavlovian motivational processes and the systems that mediate them are thought to contribute to obesity and addictions, as well as to normative reward-seeking behavior (Robinson Berridge, 2008; Dagher, 2009; Berridge et al., 2010; Volkow et al., 2013; Derman Ferrario, 2018). Thus, refining the psychological and neuronal boundaries and overlap between these mechanisms of behavioral control is critical for developing a complete understanding of the neurobiology of motivation.

Early research attempting to identify the neural circuits mediating PIT was initially hampered by the absence of a clear framework distinguishing SO-PIT, SS-PIT, and General-PIT. Although it was clear that the amygdala and the nucleus accumbens (NAc) were critical for the expression of PIT (Blundell et al., 2001; Corbit et al., 2001; Hall et al., 2001), studies examining the role of specific subnuclei produced seemingly contradictory results. Within the amygdala, lesions of the basolateral nucleus (BLA) blocked SS-PIT, but not SO-PIT; whereas in the NAc, lesions of the Core blocked SO-PIT, but not SS-PIT, yet lesions of the Shell blocked SS-PIT, but not SO PIT (Blundell et al., 2001; Corbit et al., 2001; Hall et al., 2001). Given the lack of a recognized distinction between SO- and SS-PIT, these findings were initially confusing. In an effort to clarify these discrepant findings with respect to the role of the amygdala in PIT, Corbit and Balleine (2005) designed an approach to measure General and SS-PIT within the same subject. This enabled them to evaluate the effects of BLA or central nucleus (CN) lesions on both SS- and General-PIT (Corbit Balleine, 2005).

In this procedure, rats were trained with two distinct response-outcome contingencies (R1-O1, R2-O2) and three distinct CS-Outcome associations (CS1-O1, CS2-O2, CS3-O3). Importantly, the instrumental actions shared common outcomes with CS1 and CS2, but not with CS3. With this design, SS-PIT could be measured by contrasting the effects of CS1 and CS2 on instrumental responding, whereas presentation of CS3 enabled evaluation of General-PIT, all within the same subject. This study provided the first evidence that the ability of a CS to modulate expression of instrumental behaviors can arise via at least two distinct neural mechanisms. Specifically, they found that lesions of the BLA blocked SS-PIT, but not General-PIT, whereas lesions of the CN blocked General-PIT, but not SS-PIT. Following this breakthrough, lesion and inactivation studies revealed that the expression of SS-PIT relies on the BLA and the NAc Shell, whereas expression of General-PIT relies on the CN and the NAc Core (Shiflett Balleine, 2010; Corbit Balleine, 2011). However, to this date, the explicit circuitry and the cell populations of the amygdalo-striatal pathways that mediate PIT have yet to be explicitly identified.

Here, we sought to refine the understanding of the neuronal populations involved in the expression of SS-PIT using current viral approaches to selectively reduce activity of glutamatergic neurons within the BLA during PIT testing. This was accomplished using viral-mediated expression of a Gi coupled Designer Receptor Exclusively Activated by Designer Drugs (DREADDS; hM4Di; Armbruster et al., 2007). To selectively target hM4Di expression to glutamatergic neurons we utilized a CamKII promotor which restricts expression to CamKII expressing cells. CamKII is a protein kinase whose expression has been shown to be largely restricted to glutamatergic neurons (Jones et al., 1994). Using an adapted version of the procedure pioneered by Corbit and Balleine (2005) which enables testing for both SS- and General-PIT in the same session, we determined whether activation of the Gi coupled DREADD in CamKII BLA neurons would attenuate SS-PIT, but not General-PIT. In addition, we tested the effects of CamKII BLA inhibition on the expression of Pavlovian outcome devaluation effects following conditioned taste aversion. The goal in this latter part of our studies was to determine if our DREADD manipulation alters the ability to recall a current sensory-specific representation of the CS-outcome (CS-O) relationship, in order to inform our interpretation of the nature of hM4Di-specific effects on SS-PIT. Specifically, a loss of SS-PIT can theoretically occur via disruption of the Stimulus-Outcome (S-O) association (i.e., an inability to recall the specific outcome associated with a CS) or via disruption of the Response-Outcome association (R-O, i.e., an inability to recall the specific outcome associated with a given response). This is based on the idea that SS-PIT arises when presentation of a CS activates a sensory-specific memory of the predicted outcome (CS-O), which in turn activates the instrumental R-O associative memory, thereby selectively invigorating the distinct motor response of that instrumental association (i.e., S-O-R accounts of SS-PIT; de Wit Dickinson, 2009; Alarcon Bonardi, 2016; Alarcon et al., 2018). Thus, if blockade of SS-PIT by hM4Di activation is due to a disruption of the S-O aspect of PIT, then hM4Di activation should also block the expression of Pavlovian devaluation effects. On the other hand, if hM4Di activation is disrupting the O-R (i.e., R-O) branch to prevent SS-PIT, then we would not expect to see a loss of Pavlovian devaluation effects.

## II. Materials and Methods

### Subjects

Adult Sprague Dawley rats (total N=56; male, n=28; female, n=28) purchased from Envigo (Haslett, MI) were used for the study presented here. Rats were housed in groups of two or three and maintained on a reverse light-dark schedule (12/12). All behavioral experiments were performed during the dark phase. Rats were 65 days old at the start of each experiment. All procedures were approved by The University of Michigan Institutional Animal Care and Use Committee. Additional details for all procedures and housing can be found at: https://sites.google.com/a/umich.edu/ferrario-lab-public-protocols/. All behavioral training was conducted in red light conditions.

### Viral vectors and drugs

Two CamKII dependent viral vectors were used in this study: a DREADD vector, AAV(2/10) CamKII-hM4Di-mCherry (titer, 3.83×1013 vgc/ml) and a control vector, AAV(2/10) CamKII-GFP (titer, 1×1013 vgc/ml). The hM4Di DREADD used here is a Gi coupled receptor which is activated by Clozapine-N-oxide (CNO). The control vector expressed the fluorophore GFP. The plasmids for these viral vectors were purchased from Addgene, having been deposited by Bryan Roth (pAAV-CamKIIa-EGFP: Addgene plasmid # 50469; http://n2t.net/addgene:50469; RRID:Addgene_50469; pAAVCamKIIa-hM4D(Gi)-mCherry: Addgene plasmid # 50477 http://n2t.net/addgene:50477 RRID:Addgene_50477; Armbruster et al., 2007). The virus was generated by a standard triple transfection procedure by Caroline Bass (University at Buffalo) to geierate pseudotyped AAV2/10 viral preparatiois and titered by quantitative real time PCR (Xiao et al., 1998; Gompf et al., 2015).

CNO was provided by the NIDA drug supply program. CNO solution was prepared by dissolving CNO powder in 100% dimethyl sulfoxide (DMSO) and then diluting this solution with sterile saline (0.9%) to a final concentration of 5 mg/mL CNO and 5% DMSO. CNO was administered at a 5 mg/kg (i.p.) dose for all studies. The Vehicle solution for these injections was 5% DMSO. Lithium chloride (LiCl) was used for outcome devaluation. Lithium chloride (0.3M, 63.6 mg/kg) was dissolved in sterile saline (0.9%) and saline was used as the Vehicle solution for these injections (1 ml/kg, i.p.).

### Stereotaxic surgery for viral infusion

Rats were allowed to acclimate to the vivarium for 7 days before surgeries were performed. Stereotaxic surgery was performed to deliver viral vectors into the BLA. Rats were transduced with either the CamKII-hM4Di-mCherry DREADD vector (n=35) or the CamKII-GFP control vector (n=21). Stereotaxic surgeries were conducted as previously described (Derman Ferrario, 2018). Anesthesia was induced with 5% isoflurane and maintained with 1.5-5% isoflurane. For analgesia, rats were administered Carprofen (5mg/kg, s.c.; Rimadyl) pre- and postoperatively (24 hours later). Bilateral injections of virus were made using the following coordinates: AP, −2.28mm, ML, + 5.00mm from bregma, and DV, −7.2mm from dura. A volume of 0.5*μ*l of virus was injected at a rat*μ* of 1ul/min using a microliter syringe (Hamilton, 800 series, Model 85; 26 gauges) attached to a motorized pump (Harvard, Pump 11 Elite Nanomite). Before each injection the syringe was tested to ensure proper flow and the needle was lowered to −0.5 DV below the intended DV coordinate. The needle was left in place for 5 min, then raised to the DV target site and the injection was initiated. The injection lasted for 30 sec and the needle was left in place for an additional 9.5 min and was then slowly withdrawn. Rats were left to recover for 7-10 days before food-restriction and training described below.

### Behavioral Training Chambers

Instrumental training, Pavlovian conditioning, PIT testing and devaluation testing took place in standard Med Associates operant chambers housed within sound attenuating cabinets. The front panel of each chamber contained a recessed food cup into which pellet outcomes were delivered via tubes attached to hoppers located on the exterior of the chamber. The food cup was equipped with an infrared emitter receiver unit to detect beam breaks as a measure of food cup entries. Flanking the food cup were two retractable levers and two speakers and a clicker were mounted on the rear wall. Each cabinet was equipped with red and infrared LED strips and an infrared sensitive mini camera mounted overhead (Surveilzone, CC156). Taste aversion training and consumption choice testing was conducted in rectangular plastic chambers (25.4 cm X 48.26cm X 20.32 cm) in a separate room from those housing the operant chambers.

### Instrumental Training

All behavioral training was conducted in red light conditions (Derman Ferrario, 2019). Procedures were adapted from Corbit and Balleine (2005; 2011). Following recovery from surgery, rats were food restricted to 85-90% of their ad libitum weights and maintained at this weight throughout the remainder of the study. Once reaching this target, rats were trained in 3 separate sessions to retrieve pellets from the operant chamber food cups. Three distinct pellets (45mg) were used as the outcomes (Bioserv: Unflavored #F0021: Banana #F0059: Chocolate #F0299). For a given session, 20 pellets of a given flavor were delivered into the food cup on a variable time (VT) schedule of 60 sec (range, 30-90 sec).

Table 1 depicts an outline of instrumental training, Pavlovian conditioning and PIT testing. During instrumental training rats learned two distinct response-outcome associations, where pressing on 2 distinct levers was reinforced with 2 distinct outcomes (Lever1-O1 and Lever2-O2). In the first phase of instrumental training, lever presses were reinforced on a continual reinforcement (CIF) schedule. For CRF training rats were required to reach an acquisition criterion of earning 50 pellets within less than 40 min for each lever. Lever responses, food cup entries and the time to reach these acquisition criteria were recorded.

**TABLE I:** Experimental design of PIT Training. Rats are trained in two distinct phases. In the instrumental phase, rats learn 2 sperate response-outcome (R-O) associations. In the Pavlovian phase, rats learn 3 separate CS-O associations, with 2 of the outcomes from these Pavlovian associations overlap with 2 of the outcomes from the instrumental associations. During PIT testing, rats are given continuous access to the lever, and the CSs from Pavlovian conditioning are presented intermittently to determine their influence on instrumental responding. During testing, no outcomes are delivered.

| Instrumental Training | Pavlovian Conditioning | PIT Testing |
| --- | --- | --- |
| R1-O1 | CS1-O1 | CS1: R1 v R2 |
| R2-O2 | CS2-O2 | CS2: R1 v R2 |
|  | CS3-O3 | CS3: R1 v R2 |

Next, rats were transitioned to a variable interval (VI) schedule of reinforcement. The VI scheduling was conducted as follows: the first lever response to occur following passage of a given interval resulted in delivery of 2 pellets which triggered selection and initiation of a new interval. The VIs for a given session were centered on a given time interval and the VI schedule was increased across sessions as follows: VI10 (range: 5-15 sec), VI30 (range: 15-45 sec), VI45 (range: 30-60 sec), and VI60 (range: 45-60 sec). Each schedule was trained for 2 sessions, for a total of 8 instrumental VI training sessions. Each session consisted of two 20 min periods in which each lever was trained in isolation separated by a 5 min break during which both levers were retracted (45 min total). The order in which levers were trained was counterbalanced across sessions in a double alternating pattern (e.g., first lever trained of the day: L1, L2, L2, L1, L1etc.).

### Pavlovian Conditioning

After completion of the instrumental training, rats were conditioned to three distinct CS-O associations: CS1-O1, CS2-O2, and CS3-O3 (Illustrated in Fig. 1D). Each of the pellets from food cup training were used here, two of which overlapped with the outcomes used during instrumental training (i.e., O1 and O2). The CSs used were a white noise (60dB), a tone (57dB), and a click train (20Hz); each was presented for two minutes across which four pellet outcomes were randomly delivered (VT20; range: 11-30 sec). This delivery schedule ensured that pellets were never delivered within the first 10 sec of CS presentation; this allowed us to measure anticipatory conditioned food cup approach without interference of consummatory behaviors (CS-O temporal relationship shown in Fig. 1E). Each CS-O pair was trained in isolation within a 30 min daily session. Four CS-O trials were presented in each session with a variable 5 min intertrial-interval (ITI; range: 3-7 min). Levers were retracted throughout Pavlovian conditioning, and pellet delivery was not contingent upon any behavioral response. Rats underwent 3 conditioning sessions per day, each separated by 40 min. Food cup entries were recorded throughout. RO and CS-O associations were counterbalanced across rats to ensure that each pellet flavor was evenly represented in associations with the Sensory-Specific CSs (CS1 and CS2) and the General CS (CS3).

**FIG. 1:**
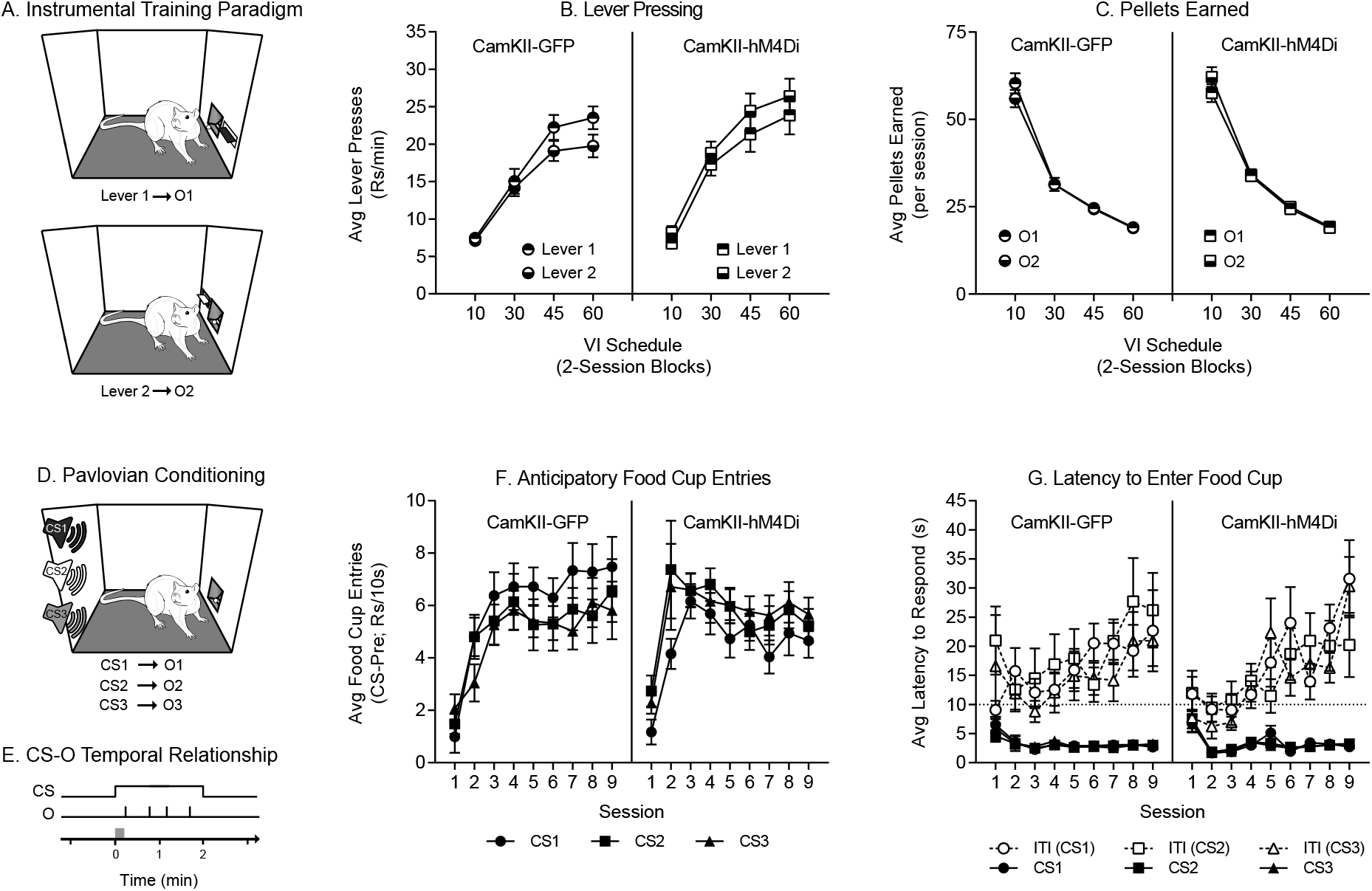
Instrumental and Pavlovian conditioning. A) Schematic of instrumental training where rats learn two independent R-O associations. B) Lever pressing on both levers increased across instrumental VI training as the schedule of reinforcement thinned. C) The number of pellets earned in VI instrumental training decreased as the schedules of reinforcement grew. D) Schematic of Pavlovian conditioning where rats were conditioned with three independent CS-O associations. E) Schematic of the temporal relationship between the CS and the paired outcomes. Each CS was presented for two minutes and four pellets were delivered randomly after the first 10 seconds following CS onset. The grey box over the time line illustrates the first 10 seconds of the CS during which pellets were never delivered. F) Anticipatory food cup entries increased within the first three sessions and then stabilized for the remaining sessions. Entries were similar across CSs. G) The latency to enter the food cup following CS onset was rapid and stable across sessions. Latencies were similar between CSs. In contrast, latencies to enter the food cup following CS offset slowed dramatically across training. All data are shown as averages ±SEM, unless otherwise noted.

### Pavlovian-to-Instrumental Transfer Testing

Rats were given an instrumental reminder session one day before each PIT test that was identical to the VI60 session described above. To determine the effect of hM4Di-mediated inhibition of CamKII BLA neurons on PIT, rats were given an injection of either Vehicle or CNO (5 mg/kg, i.p.) prior to testing. Figure 2A illustrates the injection and testing timeline. Injections were administered in the home cage where rats remained for 20 min before being placed into operant chambers for testing. Both levers were available for the entire 44 min duration of testing. Testing was conducted under extinction conditions (i.e., pellet delivery omitted). After a 10 min instrumental extinction phase, the CSs were presented in a quasi-random order separated by a fixed 2 min ITI (3 trials per CS). Lever responses and food cup entries were recorded throughout and sessions were video recorded. Rats were tested once under each treatment condition (Vehicle, CNO), with the order of treatment assignments counterbalanced across rats.

**FIG. 2:**
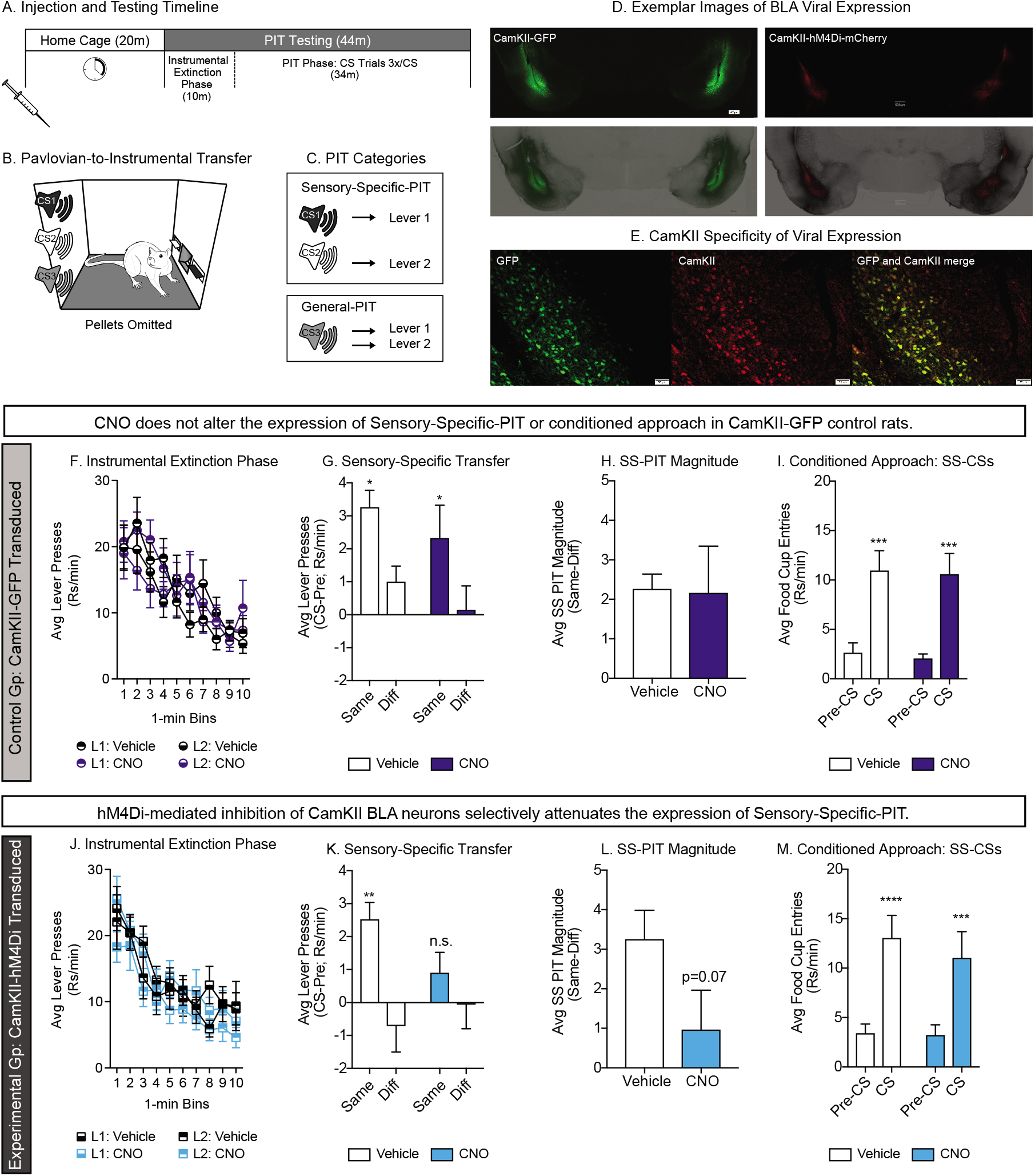
Effects of hM4Di-activation in CamKII BLA neurons on PIT. A) Timeline for testing. Rats were injected in the home-cage and then tested 20 min later. B) Schematic of PIT testing; rats were given access to both levers under extinction conditions and CSs were presented intermittently after an initial 10 min lever extinction period. C) Illustration of stimuli and responses used to measure SS- and General-PIT. D) Exemplar images of BLA CamKII-GFP (left) and CamKII-hM4Di-mCherry (right) expression. E) Exemplar image of CamKII specificity in a CamKII-GFP transduced sample tissue. F) Lever pressing in GFP transduced rats decreased across the first 10 minutes of testing, prior to CS presentation and was similar following Vehicle and CNO injections. G) SS-PIT in GFP transduced rats was unaffected by CNO administration, as is evident by greater lever pressing on the lever previously generating the Same versus the Different outcome than predicted by the CSs, following both Vehicle and CNO injections. H) In GFP transduced rats, the magnitude of SS-PIT was similar following Vehicle and CNO injections. I) In GFP transduced rats, conditioned approach was similar following Vehicle and CNO injections. J) Lever pressing in hM4Di transduced rats decreased across the first 10 minutes of testing, prior to CS presentation and was similar following Vehicle and CNO injections. K) In hM4Di transduced rats, CNO disrupted SS-PIT demonstrated by the loss of preferential responding on the Same lever following CNO injections. L) In hM4Di transduced rats, SS-PIT magnitude was diminished by CNO administration. M) In hM4Di transduced rats, conditioned approach was unaffected by CNO administration; *=p¡0.05, see results for specifics of comparisons.

### Conditioned Taste Aversion Training

To examine effects of CamKII BLA neuronal inhibition on the expression of Pavlovian outcome devaluation, a subset of rats (n=29; GFP, n=10; hM4Di, n=19) underwent conditioned taste aversion (CTA) training following the final PIT test. The purpose of CTA was to devalue one of the three outcomes from Pavlovian training (procedure adapted from Derman et al., 2018) in order to determine whether CNO disrupts the subsequent expression of Pavlovian outcome devaluation on conditioned approach. Outcome devaluation was achieved by pairing one outcome with post-ingestive injections of LiCl to induce temporary illness. For this procedure, rats were placed into individual chambers each outfitted with a metal tube feeder filled with a preweighed amount of one of the established USs (20 g, 445 pellets) and left to eat freely for 20 min (see Fig. 6.4A for schematic). Rats were then removed from these chambers and immediately injected with either saline (control sessions) or LiCl (63.6mg/kg, i.p.; devaluation sessions) and placed back in their home cages in the absence of any food. Unconsumed pellets, including spillage, were weighed to determine the amount consumed in each session. To prevent any carryover of taste aversion to their home cage lab chow, rats were fed no earlier than 2 hrs post injection. CTA training was conducted in 5, 3 session cycles. Each cycle consisted of one devaluation session and two Vehicle control sessions. Outcome devaluation assignments were counterbalanced across rats within each group.

### Devaluation Testing

To test the effect on hM4Di activation on the expression of Pavlovian devaluation, rats were injected with Vehicle or CNO 20 min prior to devaluation testing. Test sessions lasted for 36 min and were conducted under extinction conditions. Each CS was presented 3 times, in a quasi-random order, separated by a fixed 2 min ITI. Food cup entries were recorded throughout and each test session was video recorded. Rats were tested once under each treatment condition, where the order of treatment assignments was counterbalanced based on consumption during the final cycle of CTA training.

### Choice Consumption Testing for Taste A version

To test the effect of hM4Di activation on the expression of CTA, rats were given Vehicle or CNO injections 20 min prior to choice consumption testing. Each test consisted of a 20-min session in which rats were given ad libitum access to all three outcomes from training. Testing was conducted in the same chambers as initial CTA training; each chamber was outfitted with 3 feeder tubes each filled with a pre-weighed amount of one of the outcomes from training (10.2g; 227 pellets). Once testing was completed, rats were returned to their home cages and the remaining pellets were weighed. Rats were tested under both treatment conditions where the order of treatment assignments was counterbalanced based on consumption levels in the final cycle of CTA training.

### Histology and Fluorescent Immunochemistry

For brain extraction, rats were injected with a fatal dose of pentobarbital and perfused transcardially with PBS followed by 4% paraformaldehyde in PBS (PFA; w/v). The brain was then extracted, placed into a 50/50 mix of 4% PFA and 30% sucrose (w/v), and stored at 4. Approximately 24hrs later, brains were transferred to a 30% sucrose solution (2-3 days) and then sectioned. Brains were sectioned coronally at 60 microns using a cryostat (Leica) and sections were stored in cryoprotectant (50% 0.1M Phosphate Buffer; 30% Ethylene Glycol; 30% Sucrose) at −20C until being processed for immunohistochemistry (IHC). Free-floating IHC was performed to evaluate viral expression. In addition, a control study was conducted to verify the specificity of viral expression to CamKII expressing neurons. Briefly, sections were washed 12 times with 1X Phosphate Buffered Saline (PBS; 10 min/wash) and then blocked for 1.5 hrs (5% Normal Goat Serum; 0.04% Triton-X; 95% 1X PBS, room temperature). Tissue was then incubated with a primary antibody in blocking solution overnight (15-20 hrs; see below for details for each antibody used). Tissue was then washed 5 times in 1X PBS (5 min/wash) and incubated in with a secondary antibody in blocking solution for 1.25-1.5 hrs, after which it was washed again (5 times in 1X PBS, 5 min/wash) and then mounted onto Superfrost Plus microscope slides (Fisherbrand) and coverslipped with Prolong Gold +DAPI mounting medium (Invitrogen, P36931).

All IHC was conducted at room temperature using a standard orbital shaker (Talboys, NJ). All primary and secondary antibodies were incubated at a 1:2000 dilution. The following primary antibodies were used: rabbit anti-IFP (Rockland, 600-406-379); rabbit anti-GFP (Invitorgen A6455); rabbit anti-Parvalbumin (Abcam, AB11427). The secondary antibodies were used at a dilution of 1:2000: Alexa Fluor 555 Goat anti-Rabbit (Invitrogen, A32732); Alexa Fluor 555 Goat anti-Mouse (Invitrogen, 32727): Dy-Light 488 Goat anti-Rabbit (Invitrogen, 35553); Alexa Fluor 488 Goat anti-Mouse (Invitrogen, A32723).

Sections were then visualized using an upright epifluorescence manual system microscope (Olympus, BX43) with an XM10 camera; images were taken at 2x, 10x, and 20x (cellSens). Assessment of viral expression location was performed visually, using standard anatomical landmarks to identify the BLA (Paxinos Watson, 2007). For hM4Di transduced rats, only data from subjects with bilateral hM4Di-mCherry expression localized to the BLA were included for analysis (n= 20 included, 15 rejected).

### Experimental Design and Statistical Analysis

All behavioral experiments were designed for within subject comparisons. To control for unintended effects of viral transduction and potential off target effects of CNO, a viral control group in which GFP was expressed under a CamKII promoter was included in all testing. Data were processed and organized with Microsoft Excel (Version 16.16.16) and statistical analyses were performed using the GraphPad statistical software suite Prism (Version 8.0.2). Data were then assessed using, students t-tests, one-way ANOVAs, repeated measures ANOVAs (RM ANOVAs) and Holmes-Sidaks tests for planned and post-hoc multiple comparisons. Instrumental and Pavlovian behavioral data were analyzed as response rates per minute or per 10 sec and, when relevant, as a change from pre-CS rates.

For instrumental responding during VI training, the rate of responding under each VI schedule was averaged for each rat. For Pavlovian training, PIT testing, and Pavlovian devaluation testing the data were averaged across trials within a session, with the exception of lever responding in the instrumental extinction phase at the start of each PIT test. For this phase, data were analyzed in 60 sec bins.

Our goal in this study was to determine the role of CamKII BLA neurons in the expression of PIT and outcome devaluation effects on conditioned approach. As such, we set inclusion criteria to limit our analysis to subjects that exhibited the expectant behaviors under Vehicle conditions. The inclusion criterion for expression of SS-PIT was that the CS elicited greater lever responding on the lever whose outcome was the same versus the lever whose outcome was different than that predicted by the CS being presented (Same>Diff; as defined by Colwill Motzkin, 1994; Delamater Holland, 2008). For General-PIT the inclusion criterion was that CS elicited lever responding had to be greater than pre-CS lever responding (CS>Pre; averaged across levers). The inclusion criterion for rats exhibiting devaluation effects was that CS elicited food cup approach must be greater during presentation of the non-devalued CSs versus the devalued CS (N-Dev>Dev; averaged across N-Dev CSs). For post-devaluation testing, the inclusion criterion for post Devaluation testing was that CS evoked food cup approach rates to the non-devalued CSs were greater than to the devalued CS. Finally, inclusion criteria for CTA testing was that consumption of the non-devalued outcome was greater than the devalued outcome. All Ns for final groups are given in results below.

## III. RESULTS

### Histology

Exemplar images of bilateral transductions are shown in Fig. 2D for CamKII-hM4Di-mCherry and CamKII-GFP expression. IHC approaches were used to amplify mCherry or GFP expression in order to assess transduction sites. Among rats transduced with CamKII-hM4Di, 20 had bilateral on target transduction sites, 9 had bilateral transductions that were off target, and 6 had unilateral transduction. Only CamKII-hM4Di rats with bilateral, on target transduction sites were included in analyses (n=20). Among rats transduced with the control CamKII-GFP, 10 were bilateral on target, 4 were bilateral, but off target, 5 had unilateral, and 2 showed no sign of transduction. We did not find notable behavioral differences between these transduction conditions within this control group, thus data were included from all rats (n=21). Data below describing instrumental training, Pavlovian conditioning and subsequent testing include only those rats with viral expression meeting the above description.

Additional IHC was performed to qualitatively confirm that viral expression was limited to putative glutamatergic neurons (i.e., positive for CamKII and negative for parvalbumin, a marker of GABAergic interneurons). Exemplar images from this control study are shown in Fig. 2E and Supplemental Material, Fig. S1. We found nearly exclusive overlap between virally transduced cells and cells positive for CamKII labeling, and no overlap of transduced cells and parvalbumin labeled cells.

### Instrumental Training

Rats were first trained to press one lever to receive one flavored outcome (i.e., food pellet) and another lever to receive a different flavored outcome on a continual reinforcement schedule in separate sessions (Fig. 1A; Lever 1-O1 and Lever 2-O2). Rats were trained to an acquisition criterion of earning 50 consecutive pellet deliveries before moving on to a VI schedule (see also methods). The mean time to acquire this task was 24.6 min (±SEM: 3.7) and did not differ between levers within either group (Data not shown, Paired t-test, Lever 1 versus Lever 2; CamKII-GFP: p=0.60: CamKII-hM4Di: p=0.14). Next, rats were transitioned to a variable interval (VI) schedule of reinforcement that was made leaner across sessions to encourage higher rates of responding. As expected, the rate of lever pressing increased as a function of VI schedule in each group (Fig. 1B. CamKII-GFP: Mixed-effects analysis: main effect of VI schedule: *F*_(3,48)_=58.37, p<0.01: CamKII-hM4Di: Two-way RM ANOVA: *F*_(3,51)_=51.31, p<0.01). Inversely, the number of outcomes earned decreased as a function of VI schedule in each group (Fig. 1C. CamKII-GFP: Mixed-effects analysis: main effect of VI schedule: *F*_(3,48)_=163.7, p<0.01: CamKII-hM4Di: Two-way RM ANOVA: *F*_(3,51)_=437.6, p<0.01), as expected.

### Pavlovian Conditioning

Rats were next conditioned to associate 3 distinct CS-O pairs (Fig. 1D). In this procedure, outcomes were never delivered within the first 10 sec of CS presentation, thus providing a window during which true conditioned anticipatory responding could be measured across training. Fig. 1E depicts the temporal structure of the CS-O contingencies used. Anticipatory conditioned food cup approach rapidly increased across the first 3 sessions and then plateaued to asymptotic levels for the remaining sessions in both groups (Fig. 1F. Two-way RM ANOVA: CamKII-GFP: main effect of session: *F*_(8,128)_ =9.43, p<0.01; CamKII-hM4Di; Two-way RM ANOVA, *F*_(8,136)_ =5.99, p<0.01). As an additional measure of conditioning, we assessed the latency to enter the food cup following CS onset and offset (ITI). In both groups, the latency to enter the food cup following CS onset decreased across training (most notably between sessions 1 and 2), whereas the latency to enter following ITI onset increased across training (Fig. 1G. Twoway RM ANOVA: CamKII-GFP: main effect of session: CS: F(8,128)=5.08, p<0.01; ITI: *F*(8,128)=2.25, p=0.03; CamKII-hM4Di: main effect of session: CS: *F*_(8,136)_ =9.92, p<0.01; ITI: *F*_(8,136)_ =8.33, p<0.01). Thus, rats readily acquired an expectancy of reward following CS onset, with similar learning and magnitude of behavior supported by each of the three CS-O pairs.

### CNO Selectively Blocks Expression of Sensory-Specific PIT only in hM4Di-Expressing Rats

Next, rats were tested for PIT following injections of either Vehicle or CNO (within subject, treatment order counter balanced). The timeline for injections and testing is illustrated in Fig. 2A. PIT testing began with a 10 min instrumental extinction phase, followed by intermittent presentation of each CS (3 trials/CS). SS-PIT is observed when presentation of the Sensory-Specific CSs (CS1, CS2) elicits greater responding on the lever that previously generated the same outcome predicted by that CS versus the lever that generated a different outcome. General-PIT is observed when presentation of the General CS (CS3), which does not share an outcome with either lever, elicits an increase in responding on either lever above pre-CS levels (see schematic Fig. 2C). Analysis of the effects of CNO on SS-PIT and General-PIT were conducted separately, given that not all rats who showed SS-PIT also showed General-PIT under Vehicle conditions (SS-PIT: CamKII-GFP n=12/21; CamKII-hM4Di n=14/20). The data from CamKII-GFP controls are shown in Fig. 2, panels F-I, and data from the CamKII-hM4Di group is shown in Fig. 2, panels J-M.

In CamKII-GFP control rats, administration of CNO did not disrupt lever responding during the first 10 min of instrumental extinction, and as expected response rates dropped steadily across the 10 min phase (Fig. 2F. Mixed-effects analysis: no effect of drug: p=0.62; main effect of time, *F*_(9,135)_ =15.25, p<0.01; no drug x lever interaction, p=0.25). To determine whether CNO altered SS-PIT, planned comparisons were made between CS elicited lever responses made on the lever that shared the Same outcome as the CS being presented versus lever responses made on the other lever that previously produced a Different outcome. In CamKII-GFP controls, CNO administration did not disrupt SS-PIT. Specifically, following either Vehicle or CNO injection, CamKII-GFP controls showed comparable SS-PIT behavior, preferentially responding on the lever that shared the Same outcome as the CS being presented (Fig. 2G. Two-way RM ANOVA: main effect transfer, *F*_(1,12)_ =12.45, p<0.01; no effect of drug, p=0,37; no drug x transfer interaction, p=0.95; Holm-Sidak’s multiple comparisons test, Same versus Different: Vehicle: *t*_(12)_ =2.55, p=0.05; CNO: *t*_(12)_ =2.46, p=0.05). Similarly, the SS-PIT magnitude (Same[-]Diff), that is, the sensory-specificity of the transfer effect, was similar between Vehicle and CNO treatments (Fig. 2H. Paired t-test, p=0.94). Thus, in CamKII-GFP controls CNO did not affect the expression of SS-PIT.

We also evaluated potential effects of CNO on conditioned approach during SS-PIT trials (i.e., CS1 and CS2 trials). CamKII-GFP controls showed robust conditioned approach during these trials, and this did not differ following Vehicle versus CNO treatment (Fig. 2I. Twoway RM ANOVA: no effect of drug, p=0.81; main effect phase, *F*_(1,12)_ =36.97, p<0.01; no drug x phase interaction, p=0.94; Holm-Sidak’s multiple comparisons test pre-CS versus CS: Vehicle: *t*_(12)_ =4.8, p<0.01; CNO: *t*_(12)_ =4.9, p<0.01). Collectively, these data demonstrate that in the CamKII-GFP control group, CNO does not disrupt 1) lever responding generally, 2) the expression of SS-PIT, nor 3) conditioned approach.

In the DREADD-expressing CamKII-hM4Di group, CNO administration did not affect responding during the instrumental extinction phase (Fig. 2J. Three-way RM ANOVA: no main effect of drug: p=0.24; main effect of time, *F*_(9,144)_ =31.00, p<0.01; no drug by lever interaction, p=0.44). However, in contrast to controls, SS-PIT was selectively disrupted by CNO administration in the hM4Di-expressing group (Fig. 2K-L). Specifically, following Vehicle injection CS elicited lever pressing was greater on the lever that shared Same outcome as the CS being presented, versus the lever with Different outcome (Fig. 2K; Holm-Sidak’s multiple comparisons test: Vehicle: *t*_(13)_ =3.91, p<0.01;). In contrast, this preference was lost following CNO injection (Fig. 2K. Drug x transfer interaction *F*_(1,13)_ =3.79, p=0.07; CNO: Holm-Sidak’s multiple comparisons test: *t*_(13)_ =1.16, p=0.27). This effect is also apparent when we directly compared the magnitude of SS-PIT between Vehicle and CNO treatments (Fig. 2L. Paired Two-tailed, *t*_(13)_ =1.96, p=0.07). In these same rats, conditioned food cup approach elicited by the SS CSs was fully intact following CNO administration (Fig. 2M. Two-way RM ANOVA: main effect phase, *F*_(1,13)_ =47.80, p<0.01; no effect of drug, p=0.52; no drug x phase interaction, p=0.44; Holm-Sidak’s multiple comparisons test pre-CS versus CS: Vehicle: *t*_(13)_ =6.05, p<0.01; CNO: *t*_(13)_ =4.92, p<0.01). Thus, administration of CNO selectively disrupted the expression of SS-PIT in the DREADD-expressing CamKII-hM4Di group, without altering conditioned food cup approach.

### Effect of CNO on General-PIT

As stated above, analysis of the effects of CNO on General-PIT were conducted separately, given that not all rats who showed SS-PIT also showed General-PIT under Vehicle conditions (General-PIT: CamKII-GFP n=13/21; CamKII-hM4Di n=12/20). General-PIT is observed when presentation of the General CS (CS3) evokes an increase in lever responding above pre-CS rates. Administration of CNO disrupted this transfer effect in both control and experimental groups (Fig. 3). Specifically, in CamKII-GFP controls following Vehicle injection, presentation of the General CS elicited a robust increase in lever responding, this was completely absent following CNO administration in these same rats (Fig. 3A. Two-way RM ANOVA: main effect phase, *F*_(1,12)_ =19.38, p<0.01; phase x drug interaction, *F*_(1,12)_ =8.66, p=0.01; Holm-Sidak’s multiple comparisons test: Vehicle: *t*_(12)_ =5.32, p<0.01; CNO: *t*_(12)_ =1.15, p=0.27). A similar effect was observed in our hM4Di-expressing group (Fig. 3B. Two-way RM ANOVA: main effect phase, *F*_(1,11)_ =26.55, p<0.01; phase x drug interaction, *F*_(1,11)_ =5.87, p=0.03; Holm-Sidak’s multiple comparisons test: Vehicle: *t*_(11)_ =4.99, p<0.01; CNO: *t*_(12)_ =1.57, p=0.15). Comparison of the PIT magnitude between Vehicle and CNO conditions further illustrates this effect; CNO reduced the magnitude of General transfer regardless of the presence of hM4Di expression (Fig. 3C. Paired t-test, *t*_(12)_ =2.76, p=0.02; Fig. 3D. Paired t-test, *t*_(11)_ =2.47, p=0.03).

**FIG. 3:**
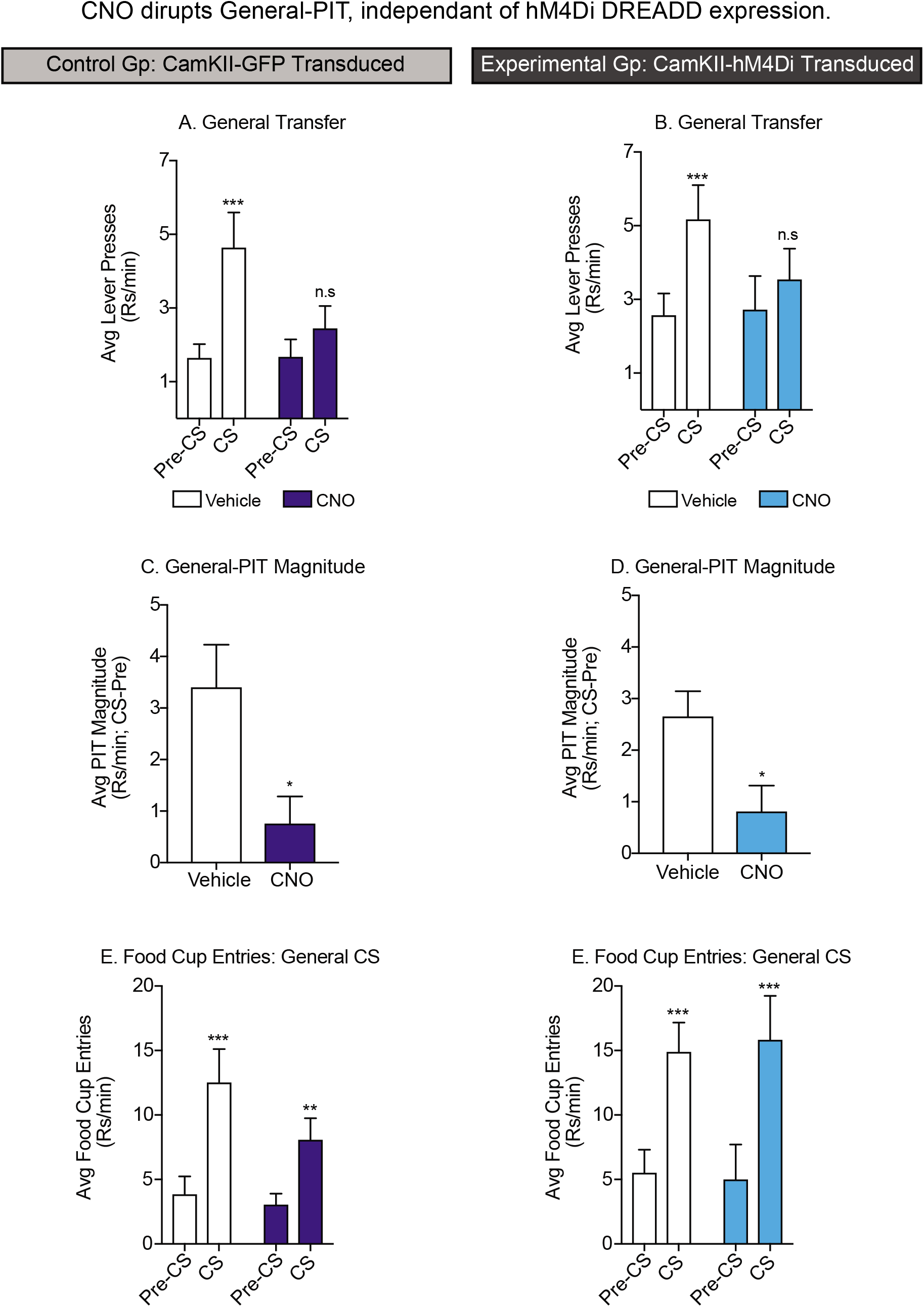
hM4Di-independent effects of CNO on General-PIT, but not conditioned approach. A) In GFP transduced control rats, CNO disrupted the expression of General-PIT. B) Similarly, in hM4Di transduced rats CNO blocked the expression of General-PIT. C) In GFP transduced control rats General-PIT magnitude is greatly diminished by CNO. D) In hM4Di transduced rats CNO reduced the magnitude of General-PIT. E) In GFP transduced control rats, conditioned food cup approach to the Gen-CS was unaffected by CNO administration. F) In hM4Di transduced rats CNO administration did not alter conditioned approach to the Gen-CS; *=p<0.05, see results for specifics of comparisons.

We also evaluated the effect of CNO on conditioned food cup approach in response to presentations of the General CS during this same testing session. Food cup entries were significantly increased above pre-CS response rates following both Vehicle and CNO injection in both the experimental and control group (Fig. 3D. Two-way RM ANOVA: main effect phase, *F*_(1,12)_ =24.94, p<0.01; no effect of drug, p=0.18; no phase x drug interaction, p=0.11; Holm-Sidak’s multiple comparisons test: Vehicle: *t*_(12)_ =5.80, p<0.01; CNO: *t*_(12)_ =3.36, p=0.01; Fig. 3E. Two-way RM ANOVA: main effect phase, *F*_(1,11)_ =60.07, p<0.01; no effect of drug, p=0.94; no phase x drug interaction, p=0.56; Holm-Sidak’s multiple comparisons test: Vehicle: *t*_(11)_ =5.47, p<0.01; CNO: *t*_(11)_ =6.31, p<0.01). Thus, although CNO disrupted the expression of General-PIT, it did not alter conditioned approach behavior. Collectively, examination of General-PIT revealed that independent of hM4Di expression, CNO exerts a robust depressive effect on General-PIT, without strongly affecting conditioned food cup approach elicited by the General CS. Therefore, CNO is not simply suppressing behavior generally or blocking recall of the CS-US association but is affecting General-PIT more specifically (see discussion).

### Conditioned Taste Aversion (CTA) Training

Following PIT testing, a subset of rats underwent CTA training. The purpose here was to devalue one of the outcomes from Pavlovian training in order to subsequently assess the effects of hM4Di activation on Pavlovian outcome devaluation effects. For each rat, one of the three outcomes from training was devalued by pairing it with post-ingestive injections of LiCl, whereas the other two outcomes were instead paired with post-ingestive saline injections. For some rats the LiCl-paired outcome was the outcome associated with the CS3 (i.e., the General CS; GFP, n=4; hM4Di, n=3), whereas for the remaining rats, the LiCl-paired outcome was one of the outcomes paired with either CS1 or CS2 (i.e., one of the SS CSs; GFP, n=6; hM4Di, n=9).

CTA data were first examined to determine if the associative nature of the devalued outcome (Dev General-O versus Dev SS-O) affected CTA acquisition. Post-ingestive pairings of the General outcome with LiCl injections (General-O: LiCl) significantly suppressed consumption of these pellets across training, whereas consumption of the SS outcome paired with saline was stable across sessions (SS-O: Sal; Fig. 4B. Two-way RM ANOVA: cycle x outcome interaction, CamKII-GFP: *F*_(8,24)_ =4.63, p<0.01; CamKII-hM4Di: *F*_(8,16)_ =6.16, p<0.01; Holm-Sidak’s multiple comparisons test: Cycle 5, SS-O1: Sal v Gen-O: LiCl: CamKII-GFP: *t*_(12)_ =5.39, p<0.01; CamKII-Hm4Di: *t*_(16)_ =8.45, p<0.01; SS-O2: Sal v Gen-O: LiCl CamKII-GFP: *t*_(12)_ =5.56, p<0.01; CamKII-Hm4Di: *t*_(16)_ =7.67, p<0.01). The same pattern was observed when the devalued outcome was one of the SS-Os and notably, there was no difference in the consumption between the non-devalued General outcome (General-O: Sal) and the non-devalued SS outcome (SS-O: Sal; Fig. 4C. Two-way RM ANOVA: cycle x outcome interaction, CamKII-GFP: *F*_(8,31)_ =14.6, p<0.01; CamKII-hM4Di: *F*_(8,64)_ =10.99, p<0.01; Holm-Sidak’s multiple comparisons test: Cycle 5, General-O: Sal v SS-O: LiCl: CamKII-GFP: *t*_(39)_ =6.09, p<0.01; CamKII-Hm4Di: *t*_(64)_ =8.61, p<0.01; SS-O: Sal v SS-O: LiCl: CamKII-GFP: *t*_(39)_ =7.69, p<0.01; CamKII-Hm4Di: *t*_(64)_ =8.32, p<0.01). Collectively, the emergence of CTA was evident by the reduction in consumption of the LiCl paired outcomes, and was present whether the devalued outcome was the General-O or an SS-O. Thus, the procedure used here reliably produced selective devaluation of the LiCl paired outcome.

**FIG. 4:**
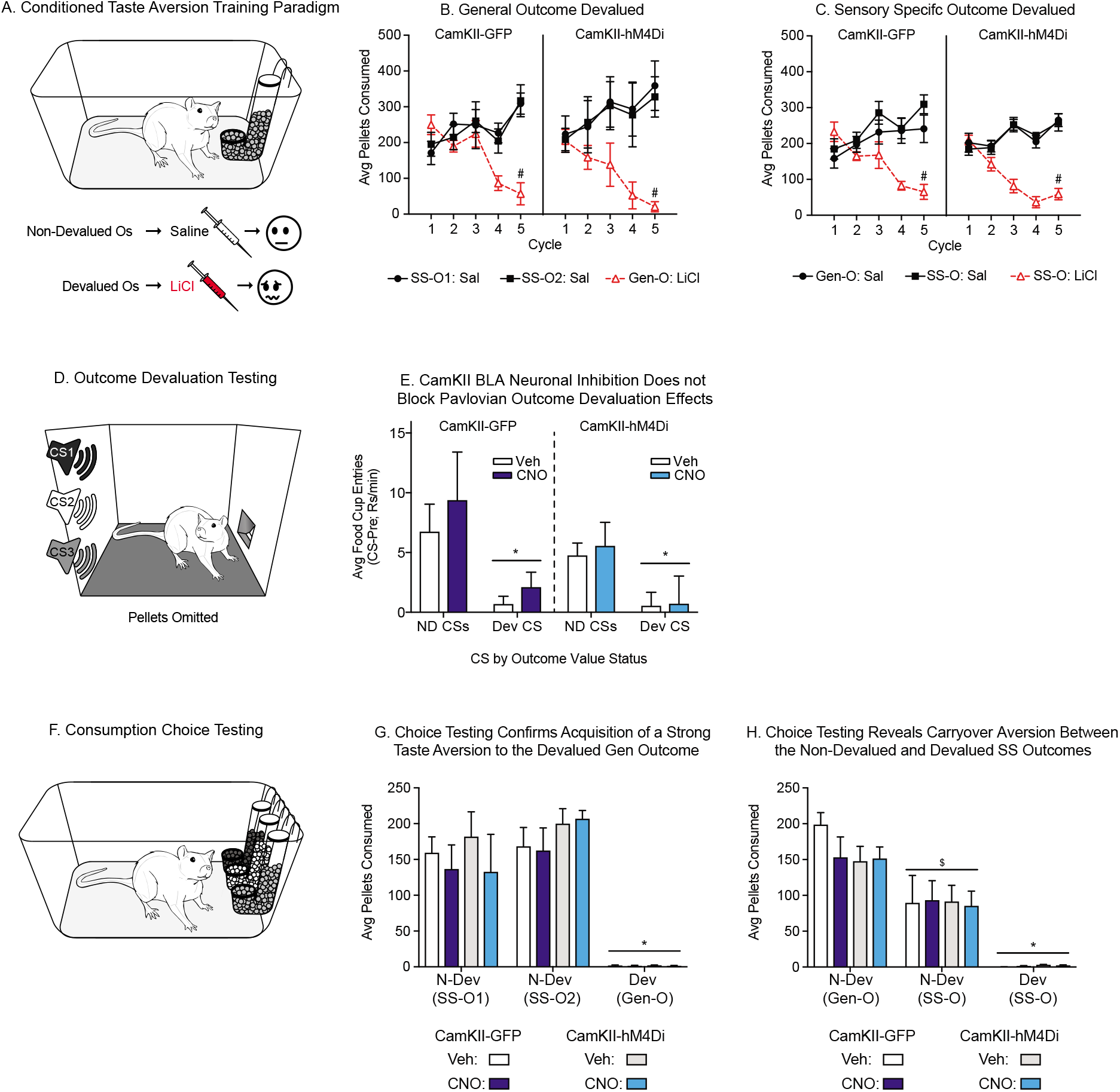
Conditioned taste aversion training, devaluation testing, and consumption choice test. A) Schematic of taste aversion training where the devalued outcome was paired with post-ingestive injections of LiCl and the non-devalued outcomes were paired with saline injections. B) Conditioned taste aversion emerged in both GFP control and hM4Di transduced rats when the LiCl paired outcome was the General outcome (#=Saline vs LiCl, p<0.05). C) Similarly, taste aversion emerges in both GFP control and hM4Di transduced rats when the LiCl paired outcome was one of the Sensory-Specific outcomes (#=Saline vs LiCl, p<0.05). D) Schematic of the test for effects of outcome devaluation on conditioned approach. E) Outcome devaluation effects were observed in rats for whom the devalued outcome was one of the Sensory-Specific outcomes. This was apparent by the reduction in conditioned approach to the presentations of the CS associated with the devalued versus the non-devalued outcomes. These effects were not altered by CNO injection in either GFP control or hM4Di transduced rats (*=NDev vs Dev, p<0.05). F) Schematic of the consumption choice test, were rats were given free access to all three outcomes in a single test session. G) All groups showed a strong conditioned aversion to the devalued General outcome with no carryover effects to the non-devalued Sensory-Specific outcomes. No differences were observed between groups, and CNO did not alter these effects. H) In contrast, conditioned taste aversion is seen to the devalued Sensory-Specific outcomes with substantial carryover effects to the non-devalued Sensory-Specific outcome. This is apparent by the strongest preference for the General non-devalued outcome over the Sensory-Specific outcome. (*=NDev vs Dev, p<0.05, $=NDev Gen-O vs NDev SS-O, p<0.05).

### Outcome Devaluation Testing

After CTA training, rats were tested for expression of Pavlovian outcome devaluation effects following Vehicle or CNO injections (within subject, treatment order counterbalanced). The purpose of this testing was to determine if hM4Di activation disrupts the ability of the CS to evoke a current representation of the outcome. Testing was performed in the operant chambers under extinction conditions (3 trials/CS). Outcome devaluation is observed when presentation of a CS whose outcome has undergone devaluation (Dev CS) evokes significantly less food cup approach than presentations of the CS whose outcome was never devalued (ND CS). This behavior depends, in part, on the ability for the rat to utilize the updated value of the outcome to appropriately guide conditioned responding and is a classic approach to examining the nature of conditioned responses, that is whether they are mediated by CS-outcome or CS-response processes (i.e., S-O, S-R; Holland and Rescorla, 1975).

Among rats for whom the devalued outcome was associated with the General CS (CS3), we did not observe reliable devaluation effects on conditioned approach following Vehicle injection in either the experimental or control groups (Supplemental Materials Fig. S2). Of 8 total rats, only 2 rats showed a reduced rate of conditioned food cup approach to presentations of the devalued CS. Thus, analysis of CNO effects on this behavior were not possible in this training group.

In contrast, the majority of rats for whom the devalued outcome was associated with one of the SS CSs expressed Pavlovian outcome devaluation effects following Vehicle injection (Total, 10/15; CamKII GFP, 4; CamKII-hM4Di, 6). Specifically, presentation of the Dev-CSs evoked fewer food cup entries than presentation of the N-Dev-CSs (Fig. 4E. Two-way RM ANOVA: CamKII-GFP: main effect of CS, *F*_(1,3)_ =6.54, p=0.08; CamKII-hM4Di: *F*_(1,5)_ =5.05, p=0.08). When CNO was given prior to testing in these same animals, the expression of this devaluation effect remained intact in both control and hM4Di-expressing groups (Fig. 4E. Two-way RM ANOVA: CamKII-GFP: no effect drug, p=0.36; CamKII-hM4Di: no effect of drug, p=0.76). Furthermore, a three-way ANOVA of viral transduction type, treatment, and CS further confirmed the expression of outcome devaluation effects in both groups following Vehicle or CNO injection (Fig. 4E. Three-way RM ANOVA: main effect of CS, *F*_(1,8)_ =11.78, p<0.01; no effect of drug, p=0.33; no drug x group x CS interaction, p=0.93). In summary, devaluation effects were not disrupted by CNO administration via either a non-specific or hM4Di-mediated mechanism among rats for whom SS-O had been devalued. This indicates that the disruption of SS-PIT observed in the hM4Di-expressing, but not control group, did not arise from an inability to retrieve a current sensory-specific representation of the outcome. Rather, the hM4Di-mediated loss of SS-PIT seems to arise from the inability to use the CS evoked memory of the outcome to preferentially enhance the appropriate instrumental response.

Additional analysis of Pavlovian devaluation effects revealed that there was carryover between CSs trained as Sensory-Specific stimuli, such that devaluation of one of the SS-CSs resulted in reduced conditioned approach elicited by the non-devalued SS-CS as compared to the non-devalued General CS. Given that these data are not directly relevant to the primary question addressed here (i.e., how hM4Di activation in BLA CamKII neurons affects the expression of SS-vs General-PIT), results and discussion of carryover effects are presented in Supplemental Materials. Importantly, these effects do not alter the interpretation of effects of CNO on SS- or General-PIT.

### Choice Consumption Test

Finally, to more stringently evaluate the efficacy of CTA and to determine whether CNO effects the expression of CTA, we performed a free choice consumption test in which all three outcomes were available. Data were analyzed separately for rats in the two different CTA training conditions (Dev Gen-O training versus Dev SS-O training) because we observed differences in expression of Pavlovian devaluation effects in these groups (see above).

In the group for which the devalued outcome was the General-O (O3), choice testing revealed a robust aversion to the devalued outcome. Consumption of both nondevalued SS-Os was significantly higher than consumption of the devalued General-O (Fig. 4G. Three-way RM ANOVA, main effect of outcome, *F*_(2,10)_ =39.38, p<0.01; Holm-Sidak’s multiple comparisons test: ND SS-O1 versus Dev Gen-O, *t*_(10)_ =6.86, p<0.01; ND O2 versus Dev Gen-O, *t*_(10)_ =8.31, p<0.01). Moreover, consumption was similar between the non-devalued outcomes (Holm-Sidak’s multiple comparisons test: ND SS-O1 versus ND SS-O2, p=0.18). Finally, this taste aversion did not differ between CamKII-GFP and CamKII-hM4Di groups, and there was no difference in this effect following Vehicle or CNO injections (Fig. 4G. Three-way RM ANOVA, no main effect of group, p=0.41; no main effect of drug, p=0.23; no drug x group x outcome interaction, p=0.80). Together these data confirm that CTA training produced the intended taste aversion, that CNO has no effect on the expression of this consummatory behavior, and that the absence of devaluation effects on conditioned approach elicited by the General CS (CS3; Supplemental Materials Fig. S2), are not due to a failure to acquire CTA (Fig. 4G).

The pattern of consumption was somewhat different in the groups for which the devalued outcome was one of the SS-Os. First, as expected, we observed a strong aversion to the devalued SS-O; rats barely consumed the devalued outcome and consumed substantially more of both non-devalued outcomes (Fig. 4H. Three-way RM ANOVA, main effect of outcome, *F*_(2,24)_=61.77, p<0.01: Holm-Sidak’s multiple comparisons test: ND Gen-O versus Dev SS-O, *t*_(24)_=11.10, p<0.01: ND SS-O versus Dev SS-O, *t*_(24)_=6.07, p<0.01). However, we observed substantial carryover of CTA of the devalued SS-O to the non-devalued SS-O: this is a pattern similar to that found for Pavlovian devaluation effects mentioned above. This was evident by substantially reduced consumption of the non-devalued SS-O compared to the non-devalued General-O (Fig. 4H: Holm-Sidak’s multiple comparisons test: ND SS-O versus ND Gen-O, *t*_(24)_=5.03, p<0.01). Importantly, these effects were not different between CamKII-GFP and CamKII-hM4Di groups and CNO did not alter this effect (Fig. 4H: Three-way RM ANOVA, no main effect of group, p=0.28: no main effect of drug, p=0.24: no drug x group x outcome interaction, p=0.67). Critically, the flavors of the assigned outcomes (banana, chocolate, or unflavored pellets) were counterbalanced across rats and groups, precluding the interpretation that similarities between the intrinsic sensory properties of the SS-Os may account for this effect.

Collectively, these data demonstrate that CNO does not attenuate the expression of CTA either through a nonspecific effect or via a hM4Di-mediated effect. Furthermore, there are carryover effects on CTA between devalued and non-devalued outcomes previously trained as SS stimuli. This final point is relevant to the utilization of this procedure more broadly, but does not impact overall interpretation of PIT results here (see Supplemental Materials for additional discussion of carryover effects).

## IV. DISCUSSION

Pavlovian conditioned stimuli often acquire the ability to control instrumental behaviors, a phenomenon known as Pavlovian-to-Instrumental transfer that can be observed in rodents (Walker, 1942; Cartoni et al., 2016) as well as humans (Colagiuri Lovibond, 2015: De Tommaso et al., 2018: Hogarth et al., 2018). This phenomenon is thought to play an important role in a wide range of behaviors that are essential for survival, and in the development of problematic appetitive behaviors that drive addictions and obesity (Wyvell Berridge, 2000; Berridge Iobinson, 2003; Bouton, 2011; Boutelle Bouton, 2015; Alonso-Caraballo et al., 2018; Derman Ferrario, 2018; Watson et al., 2018). Consistent with the preclinical literature, studies in humans also find support for alterations in PIT and its underlying neural and psychological processes in obesity and internet gaming disorders (Lehner et al., 2017; Vogel et al., 2018). Thus, understanding the neural basis of PIT is critical for addressing normal and aberrant motivation.

### A. CamKII-BLA neurons mediate the expression of Sensory-Specific PIT

In the current study we used viral mediated expression of an hM4Di DREADD under the control of the CamKII promotor to determine whether activation of hM4Di DREADDs in CamKII neurons of the BLA is sufficient to block the expression of SS-but not General-PIT. A CamKII promoter was used to target glutamatergic neurons (Jones et al., 1994). As expected, labeling of CamKII revealed that GFP expression was limited to CamKII positive cells in CamKII-GFP-AAV transduced tissue (Fig. 2E). In addition, there was no overlap between GFP expression and Parvalbumin positive cells (Sup. Fig. 1). This same pattern of results was observed with CamKII-hM4Di-mCherry transduced tissue, where mCherry expression was limited to CamKII positive cells and did not overlap with Parvalbumin cells. Thus, our viral manipulation was successful in targeting the expression of hM4Di to glutamatergic neurons within the BLA.

Effects of CNO on behavior were evaluated in CamKII-hM4Di-mCherry transduced and CamKII-GFP transduced groups, allowing us to control for viral transduction and to examine potential effects of CNO alone (e.g., Gomez et al., 2017). We found that CNO blocked the expression of SS-PIT in the hM4Di-expressing group, but not in GFP-expressing controls (Figs 2G, K). This is consistent with previous lesion and inactivation studies (Corbit Balleine, 2005; Shiflett Balleine, 2010) and identifies CamKII expressing BLA neurons as key mediators of SS-PIT. Furthermore, CNO administration did not block the expression of conditioned approach during PIT testing (Figs 2I M; Fig. 3E, F). This is important because it shows that effects on SS-PIT are not due to a disruption in the ability to recall the reward-predictive nature of the CS. Finally, Pavlovian outcome devaluation effects were also intact following hM4Di activation in control and experimental groups (Fig. 4E), indicating that the effect of hM4Di activation on SS-PIT was not the result of an inability to recall the sensory-specific representation of a given outcome evoked by a CS. Thus, hM4Di-mediated inhibition of CamKII BLA neurons prevented the CS-O representation from initiating the appropriate instrumental response (de Wit Dickinson, 2009: Alarcon Bonardi, 2016; Alarcon et al., 2018). This interpretation is further supported by examination of Pavlovian outcome devaluation effects discussed below.

### B. hM4Di-mediated loss of SS-PIT is not due to a disruption of the sensory-specific CS-O representation

Following PIT testing, a subset of rats underwent CTA followed by testing for expression of Pavlovian outcome devaluation effects after CNO or Vehicle injection. The purpose of this experiment was to determine whether hM4Di-mediated attenuation of SS-PIT was the result of an inability for the CS to call up a current sensory-specific representation of the outcome, one of the primary mechanisms by which SS-PIT is thought to be mediated (de Wit Dickinson, 2009; Alarcon Bonardi, 2016; Alarcon et al., 2018). We found that CNO administration did not alter expression of Pavlovian outcome devaluation effects on conditioned approach (Fig. 4E). This is consistent with previous studies suggesting that disruption of BLA function at the time of testing does not alter the expression of Pavlovian devaluation effects (Blundell et al., 2003; Pickens et al., 2003; Wellman et al., 2005); see below for additional discussion of this point. Thus, the absence of effects of CNO on Pavlovian outcome devaluation strongly suggest that CNO-induced disruption of SS-PIT in the CamKII hM4Di-expressing group does not arise from an inability to call up the sensory-specific details of the CS-O associations. Rather, the effect of hM4Di activation is likely due to an inability of the CS-evoked memory of the outcome to access the appropriate motor networks mediating the instrumental transfer effect. In other words, assuming an S-O-R framework of PIT, BLA CamKII neurons appear to be critical for the ability of the outcome memory to activate the appropriate instrumental memory.

Finally, we also evaluated the efficacy of our CTA manipulation and potential effects of CNO or hM4Di activation on sensory processing more generally. When given a free choice between previously devalued and non-devalued outcomes, rats exhibited robust avoidance of the devalued outcome following Vehicle injection. Thus, our procedure induced strong avoidance of the LiCl paired outcome. Furthermore, CNO injection did not alter this avoidance in GFP control or hM4Di-expressing groups (Fig. 4G). This confirms that neither hM4Di activation in CamKII BLA neurons, nor CNO itself disrupt sensory processing of the outcome or the ability to recall the recently updated post-ingestive effects of these outcomes. These results are consistent with previous lesion and inactivation studies demonstrating that expression of CTA occurs independent of the BLA (Blundell et al., 2003; Pickens et al., 2003; Wellman et al., 2005), and shows that CNO administration itself does not produce generalized effects on consummatory behaviors.

### C. CamKII-BLA neurons do not mediate the expression of Pavlovian outcome devaluation effects

Although not the primary focus here, our data show that CNO does not affect the expression of Pavlovian devaluation effects in GFP control or hM4Di-expressing groups (Fig. 4E). As mentioned above this is consistent with some previous studies. However, it is worth noting that evidence for the necessity of the BLA in the expression of Pavlovian outcome devaluation effects is mixed. Some studies have supported its role (Baxter et al., 2000; Johnson et al., 2009; Lichtenberg et al., 2017) and yet others mentioned above, (Blundell et al., 2003; Pickens et al., 2003; Wellman et al., 2005), and our current results, show that disruption of the BLA at the time of testing does not alter the expression of Pavlovian devaluation effects. Among these studies, only two conducted manipulations of the BLA at the time of testing, as we did here (Wellman et al., 2005; Lichtenberg et al., 2017). Again, one study finding evidence for involvement of the BLA, and the other not. However, the discrepant results in these cases may be more readily explained by differences in the depth of devaluation and in the sensory-specificity of the initial CS-O training. For example, Lichtenberg et al., (2017) found that inhibition of BLA to orbital frontal cortex (OFC) efferents at the time of testing blocked Pavlovian outcome devaluation effects in rats using a 1-hour pre-feeding satiety induced devaluation procedure. Wellman et al., (2005) found that muscimol induced BLA inactivation at the time of testing did not disrupt satiety induced Pavlovian outcome devaluation effects in monkeys. In their study, monkeys had ad libitum access to the outcome, and testing was only performed once each subject had stopped consuming the food, whereas in Lichtenberg et al., (2017) the pre-feeding period was fixed and satiety was not confirmed via consumption testing. Potential species differences aside, results here and in the Wellman et al., (2005) study both found no effect of BLA manipulations at the time of testing on Pavlovian outcome devaluation effects, and both used procedures that produced clear and pronounced outcome devaluation. Thus, apparent discrepancies in determining the necessity of the BLA for expression of Pavlovian outcome devaluation effects may be explained by the degree of devaluation. This would suggest that sufficiently strong devaluation reduces the role of the BLA in the expression of this behavior.

### D. Which target nuclei of CamKII BLA efferents may mediate the expression of Sensory-Specific-PIT?

Results above expand upon our understanding of the neuronal circuitry underlying SS-PIT by identifying glutamatergic, CamKII-expressing BLA neurons as critical for the expression of this behavior. An outstanding question is which target nuclei of these CamKII BLA efferents mediate the expression of SS-PIT? Of the major efferents of the BLA (Sah et al., 2003), there is evidence for involvement of direct projections to the striatum and the OFC in the expression of SS-PIT. As mentioned in the introduction, lesions of the NAc Shell block the expression of SS-PIT (Corbit Balleine, 2011), implicating it as a likely target. However, a recent study by Lichtenberg et al., (2017) demonstrated that projections from the BLA to the OFC are also critical for the expression of SS-PIT. In this study, optogenetic inhibition of BLA efferent terminals within the OFC blocked expression of SS-PIT. Taken with previous results, it is possible that expression of SS-PIT may rely on multiple separate, and partially redundant neural circuits. Alternatively, BLA to NAc Shell versus BLA to OFC pathways may influence different aspects of the PIT phenomenon. Studies examining the contribution of each of these circuits to the expression of SS-PIT within the same subject would help address these possibilities.

Of course, other cell populations within the BLA may also influence its output. For example, cholinergic neurons indirectly influence activity of glutamatergic output neurons in the BLA (Lang Pare, 1998; Woodruff Sah, 2007; Spampanato et al., 2011; see Prager et al., 2016 for review). Thus, it is likely that perturbations of local circuits within the BLA, by either directly targeting BLA GABAergic interneurons or targeting their cholinergic afferents, may also alter the expression SS-PIT.

### E. CNO disrupts General-PIT, in the absence of hM4Di expression

Interestingly, while effects of CNO on SS-PIT were selective to the hM4Di-expressing group, General-PIT was reduced by CNO in both the hM4Di-expressing experimental group and GFP-expressing controls (Fig. 3A-D). However, in these same rats CNO did not disrupt conditioned approach or responding during the initial 10 min instrumental extinction phase of the test (Fig. 3E, F). Similar null effects of CNO during extinction and on conditioned approach were seen in the hDM4Di transduced group (Fig. 2 F,I). In addition, CNO also had no effects on Pavlovian devaluations effects (Fig. 4E) or food consumption during free choice testing (Fig. 4G-H). Thus, while we did find an hM4Di independent effect of CNO it was still specific in nature, blocking the expression of General-PIT, but not any other Pavlovian responses or motor performance per se. We suspect that this specific effect of CNO on General-PIT may be due to the particular sensitivity of General-PIT to internal state, addressed in the following section.

One of the interesting distinctions between SS-PIT and General-PIT is that the expression of General-PIT can be altered by shifts in internal state, whereas SS-PIT is stable across states. For example, testing in a satiated (rather than hungry) state abolishes the expression General-PIT, but does not block the expression of SS-PIT (Corbit et al., 2007; though see Watson et al., 2014). Similar results have been observed with thirst (Balleine, 1994; De Tommaso et al., 2018). There is no evidence that CNO alters hunger or thirst, but a recent study demonstrated that CNO can, in fact, produce a detectable state in rats that may be related to the metabolism of CNO into the psychoactive compound Clozapine (Gomez et al., 2017; Manvich et al., 2018). Specifically, Manvich et al., (2018) first trained systemic Clozapine (1.25 mg/kg) versus saline injection as a discriminative stimulus during an instrumental task, such that one lever was reinforced under Saline conditions, whereas a different lever was reinforced under Clozapine conditions. After establishing this discrimination, rats were challenged with CNO (1.0, 3.2, 10 mg/kg, i.p.) and lever preference was tested under extinction conditions. During testing, rats injected with 10 mg/kg CNO (but not 1.0 or 3.2 mg/kg) preferentially responded on the lever previously reinforced following Clozapine injection. Thus, administration of 10 mg/kg CNO produced a state that was distinguishable from saline, and may share features of the state induced by Clozapine. In the current study, we used a 5 mg/kg dose of CNO. Thus, it is possible that the disruption of General-PIT, observed in GFP-expressing controls here may be due to a shift in internal state that disrupted the disrupted General-but not SS-PIT. Taken as a whole our data show that its possible for the same dose ofCNO to have hM4Di DREADD-mediated AND DREADD independent effects, depending on the behavior being examined.

To our knowledge, no studies have directly tested whether drug-induced shifts in state have a similar effect on General-PIT as shifts in thirst or hunger. However, given the labile nature of General-PIT, that we observed a loss of General-PIT following CNO administration, and that CNO can produce a state that is dissociable from saline (Manvich et al., 2018), it is likely that General-PIT may be shifted by a wider range of states than previously identified. This also opens the possibility of using drug-induced state shifts to alter undesired cue-triggered motivation that arises via general affective processes. Finally, these results also suggest that the expression of General-PIT may be an alternative paradigm for studying the broader affective properties of psychoactive drugs. Studies addressing this possibility are underway.

Regarding the use of DREADDs and CNO more broadly, these results also suggest a potential challenge for researchers using this approach to examine circuits and cell populations mediating general affective motivation, as they may be more sensitive to disruption by CNO alone. Of course inclusion of appropriate control groups helps mitigate potential false positives. In addition, the use of Compound 21, which activates hM3Dq and is not metabolized into a known psychoactive compound (Chen et al., 2015), may be beneficial. However, its efficacy at the hM4Di DREADD has yet to be tested, and little is known about Compound 21’s potential behavioral effects. That CNO blocked the expression of SS-PIT only in hM4Di-expressing rats, but had hM4Di-independent effects on the expression of General-PIT highlights the need to consider both psychological and neurobiological aspects of behavior when designing and interpreting studies of this kind.

### Summary

In sum, data presented in this study refine our understanding of the neural circuits involved in the expression of PIT by showing that glutamatergic neurons within the BLA are critical for the expression of SS-PIT, but not General-PIT. In addition, effects of CNO alone on General-PIT suggest that drug induced states may also affect the expression of this behavior in a manner similar to states related to hunger and thirst. This also speaks to potential confounds of using CNO to understand the neural mechanism of affective motivation. Finally, control studies using CTA further support the specific role of BLA in SS-PIT, and also provide additional new insights into the independence of associations encoded via sensory-specific versus general affective processes. Future studies will need to identify the critical afferent sites for BLA CamKII neurons in the expression of PIT. In addition, the distinct cell populations and circuitry mediating General-PIT still remains to be elucidated.

## Acknowledgments

This work was supported by the National Institutes of Health [NIDDK 1F31-DK111194-01 awarded to ICD, I21DA043190 and Whitehall Foundation Grant 2017-12-98 awarded to CEB, and R01DK106188, R01DK115526 awarded to CIF]. Viruses plasmids were obtained from Addgene as gifts from Bryan Roth (Addgene plasmid # 50469; http://n2t.net/addgene:50469; RRID:Addgene_50469; Addgene plasmid # 50477; http://n2t.net/addgene:50477; RRID:Addgene_50477). CNO was provided by the Drug Supply program of the National Institute on Drug Abuse.

## Author Contributions

RCD designed and executed the experiments, analyzed data, and wrote the manuscript. BEC generated the viruses and wrote the manuscript. CRF designed experiments, analyzed data, and wrote the manuscript. All authors have read and approve the manuscript.

## VI. SUPPLEMENTAL MATERIALS

**FIG. S1:**
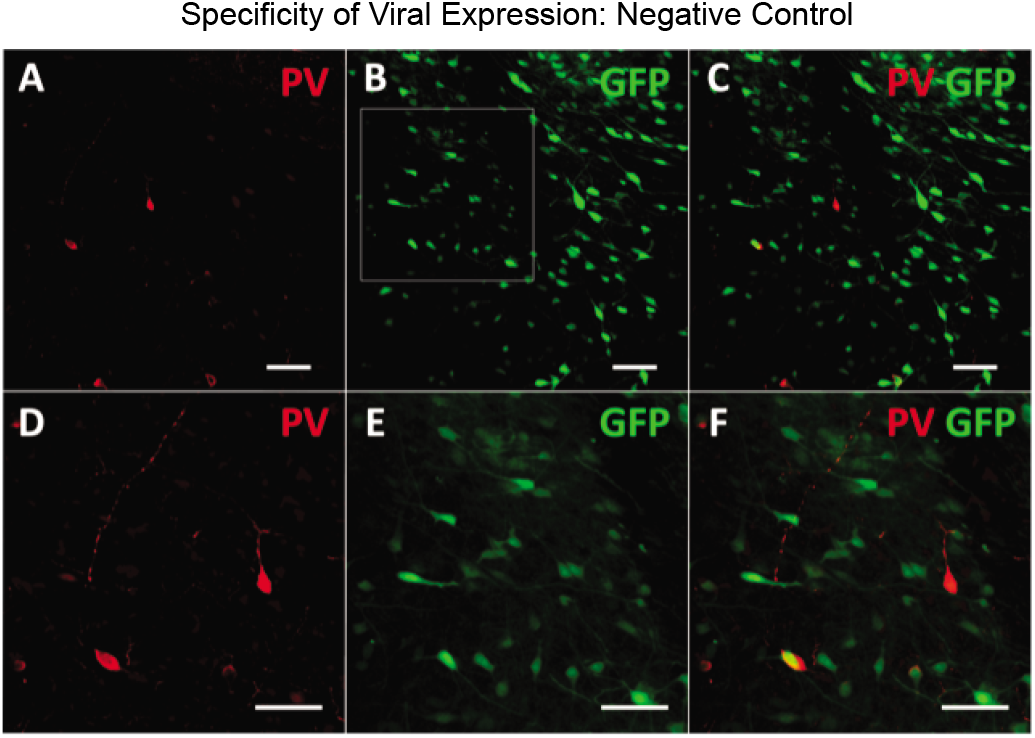
No overlap between Parvalbumin positive cells and cells transduced with CamKII-GFP. A) Parvalbumin positive (PV+) cells in the BLA (20X). B) GFP expressing cells transduced with CamKII-GFP (20X). C) Merged image of PV+ and GFP expression cells (20X). D) Close-up of PV+ cells in panel A. E) Close-up of GFP expressing cells in panel B. F) Close-up of merged image in panel C. Scale-bar in all images is 50*μ*m.

**FIG. S2:**
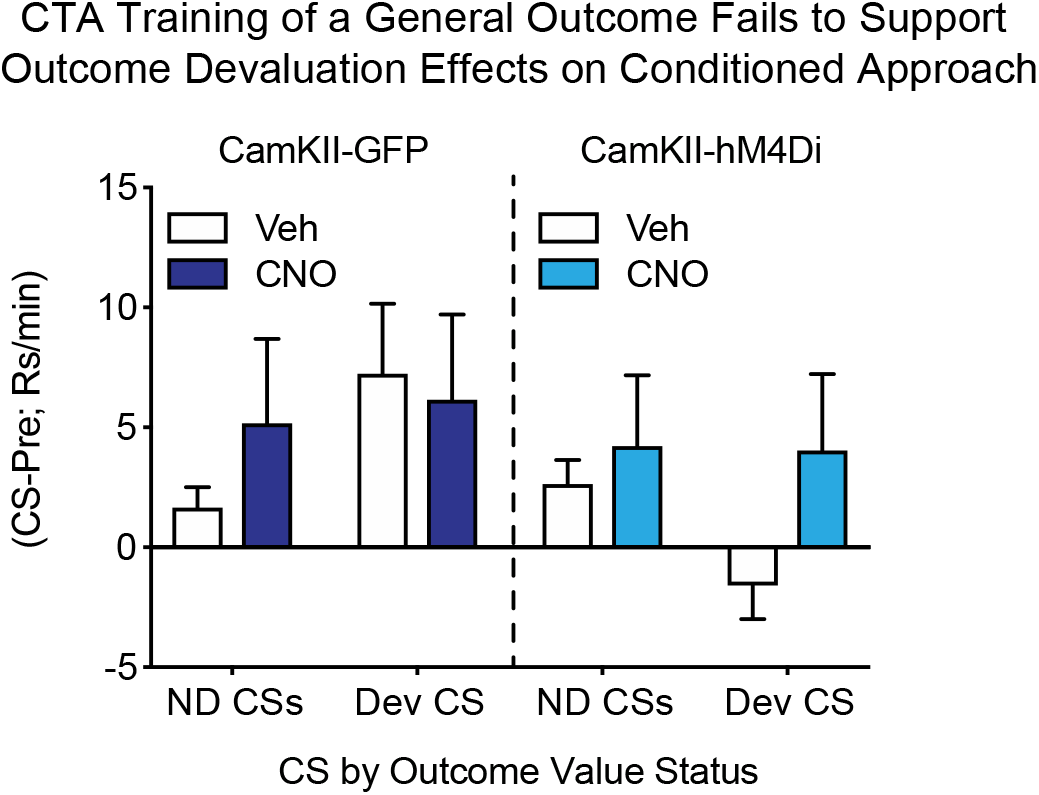
Pavlovian devaluation testing in rats where the devalued outcome was one of the previous SS outcomes (i.e, O1 or O2; Table 1). Both CamKII-GFP and the CamKII-hM4Di groups exhibited Pavlovian devaluation effects, approaching the food cup less during presentation of the devalued CS (Dev SS) vs the non-devalued CSs (NDev Gen and NDev SS). CNO administration did not alter this behavior.

### A. Examining Carryover Effects of CTA on Pavlovian Conditioned Approach

#### Results

Although not directly related to our initial hypotheses, we conducted additional analyses of the behavioral responses to CSs corresponding to the non-devalued outcomes in order to determine whether devaluation of the SS-CSs carried over to the other non-devalued SS-CS, or non-devalued Gen-CS. First, we determined whether the pattern of conditioned approach responses to these CSs (ND Gen-CS, ND SS-CS, and Dev SS-CS) differed between Vehicle and CNO conditions, and found no effect of drug in either group (Fig. S3. Three-way RM ANOVA, no effect of drug, p=0.34; no drug x group interaction, p=0.55). We then determined whether conditioned approach differed across the CSs during devaluation testing. We found that the magnitude of conditioned approach differed across each CS regardless of viral transduction or CNO treatment (Fig. S3. Three-way RM ANOVA, main effect of CS, *F*_(1,8)_ =11.78, p<0.01; no CS by group interaction, p=0.53; no CS x drug interaction, p=0.76). Therefore, we collapsed the data across drug and viral transduction conditions (Fig. S4) in order to examine conditioned approach to CS previously paired with devalued or non-devalued outcomes (ND Gen-CS, ND SS-CS, and Dev SS-CS). This revealed a strong carryover devaluation effect on conditioned approach elicited by the ND Gen-CS and the Dev SS-CS (Fig. S4, One-way RM ANOVA, main effect of CS *F*_(1.44,12.94)_ =5.46, p=0.02; Holm-Sidak’s multiple comparisons test: *t*_(9)_ =3.00, p=0.03). In contrast a weaker devaluation effect was observed between the ND SS-CS and the Dev SS-CS (Fig. S4, Holm-Sidak’s multiple comparisons test: *t*_(9)_ =2.10, p=0.07). These data demonstrate carryover devaluation effects between the SS-CSs suggesting that the shared associative properties of these outcomes, renders them similar enough to support these carryover effects.

**FIG. S3:**
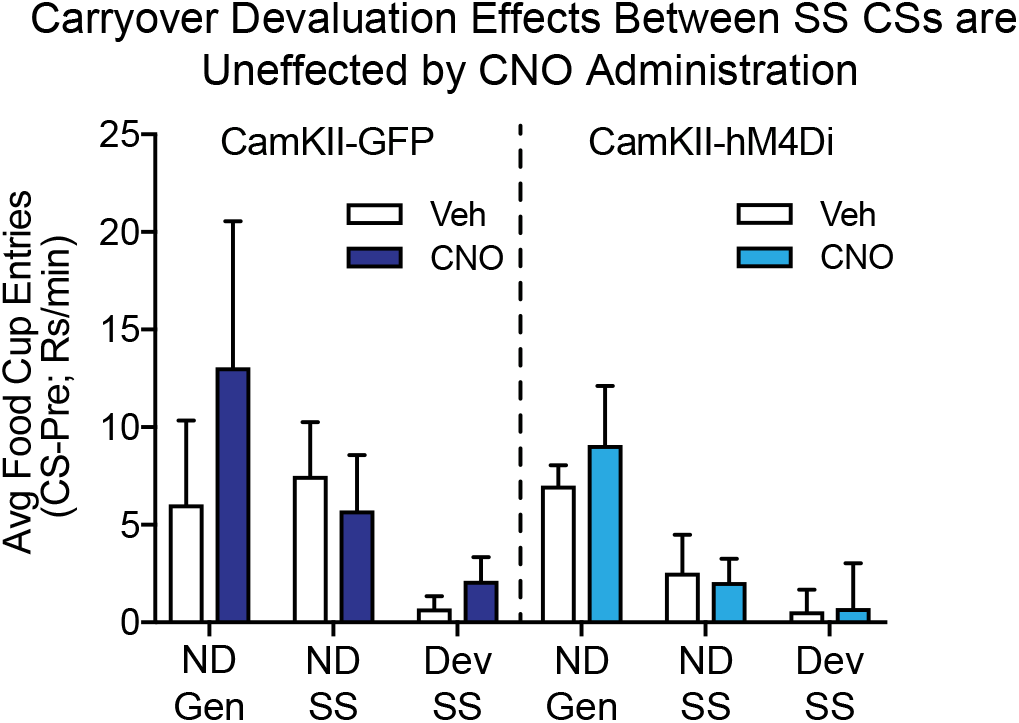
Pavlovian devaluation testing in rats where the devalued outcome was one of the previous SS outcomes (i.e, O1 or O2; Table 1). Both CamKII-GFP and the CamKII-hM4Di groups exhibited Pavlovian devaluation effects, approaching the food cup less during presentation of the devalued CS (Dev SS) vs the non-devalued CSs (NDev Gen and NDev SS). CNO administration did not alter this behavior.

#### Discussion

One interesting and unexpected behavioral finding in this study that is separate from the question of the role of BLA in PIT, was that rats showed carryover effects between outcomes trained under conditions promoting sensory-specific encoding versus general affective encoding. Specifically, we found that when CTA was performed using one of the outcomes from the CS-Os used to promote SS-PIT (e.g., O1 or O2), Pavlovian devaluation effects were seen to both the devalued SS-CS, and to the non-devalued SS-CS, but not to the non-devalued Gen-CS (Fig. S4). This carryover effect while mild, was notable. Furthermore, a conceptually similar but stronger carryover effect was observed during the taste aversion choice testing (Fig. 4H). During choice testing, rats vastly preferred the non-devalued Gen-O versus the non-devalued SS-O. Critically, the flavors had been carefully counterbalanced and therefore similarity in sensory perception of outcomes alone cannot account for this effect.

**FIG. S4:**
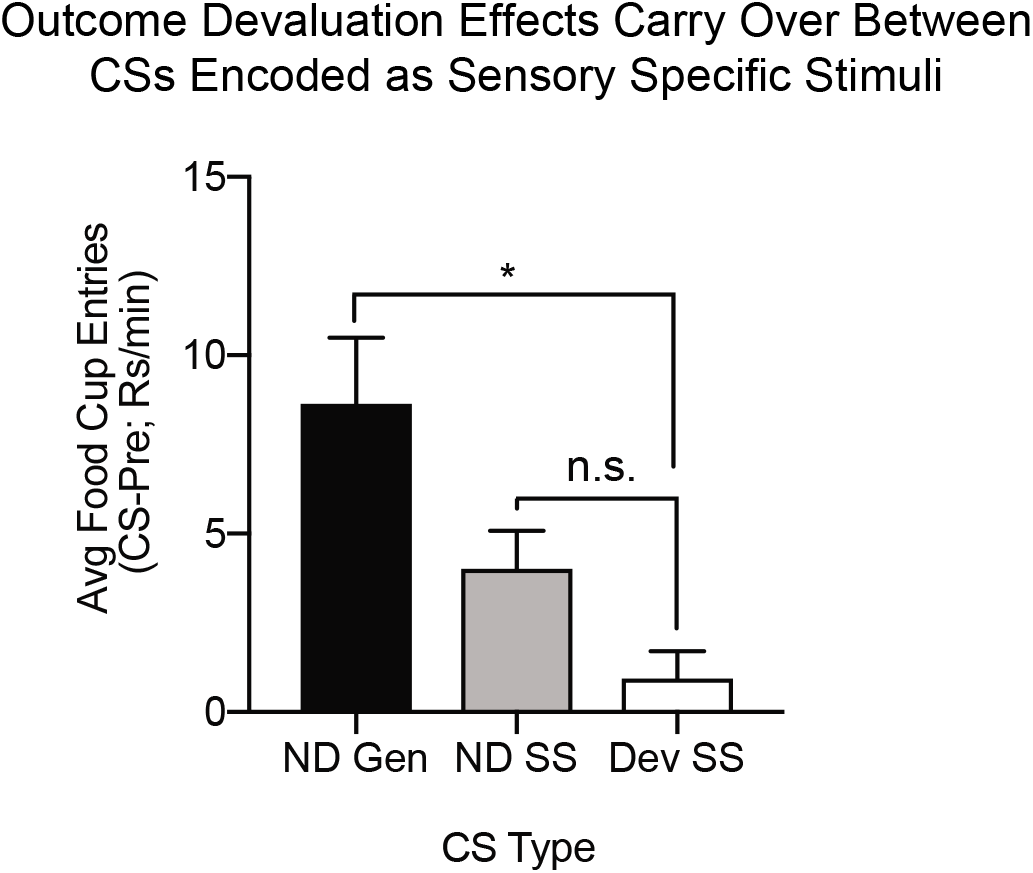
Collapsed CamKII-GFP and CamKII-hM4Di group data from Pavlovian devaluation testing in rats where the devalued outcome was one of the SS outcome (i.e, O1 or O2; Table 1). Conditioned approach to the non-devalued SS-CS is lower than to the non-devalued General-CS (NDev Gen), illustrating carryover devaluation between the devalued SS-CS (Dev SS) and the non-devalued SS-CS (NDev SS).

These data suggest that CTA can produce aversions not just to stimuli directly associated with the devalued outcomes, but also to other stimuli (outcomes and CSs) that share the same associative modalities (i.e., SS versus Gen). In this PIT paradigm, the outcomes that support SS-PIT are each independently associated with a Pavlovian CS (CS-O) and an instrumental response I-O association. In this way the outcome is embedded in a wider S-O-I network. In contrast, the outcome that supports General-PIT is only experiences within a Pavlovian CS-O association, but not an instrumental I-O association. In this way the General-O exists in a smaller S-O network. The SS-Os share a common associative structure, which is distinct from that of the General-O, which render the outcomes trained under conditions promoting SS encoding vulnerable to effects observed. If outcomes trained within the same associative history are more vulnerable to carryover effects of CTA training, then we would expect a similar effect between a devalued General-O and a non-devalued General-O, where a non-devalued SS in this paradigm should be immune to carryover. This hypothesis has yet to be tested. Nevertheless, the finding here of carryover between a devalued SS-O and a non-devalued SS-O, but not a General-O, provides further support for the idea that independent encoding processes exist for sensory specific associative structures and general affective associative structures (as canonically proposed by Konorski, 1967), and presumably dissociable neural mechanisms underlying these processes.

